# Limitations of bacterial culture, viral PCR, and tulathromycin susceptibility test methods from upper respiratory tract samples in predicting the outcome of tulathromycin control or treatment of bovine respiratory disease in high risk feeder heifers

**DOI:** 10.1101/2021.02.04.429716

**Authors:** Jeffrey J. Sarchet, John H. Pollreisz, David T. Bechtol, Mitch Blanding, Roger L. Saltman, Patrick C. Taube

**Author notes:** Corresponding author: Dr. Jeff Sarchet, Address: 123 Private Road 4011, Decatur, TX 76234.

## Abstract

A cross-sectional prospective cohort study to correlate BRD clinical outcomes for tulathromycin metaphylaxis/treatment for bovine respiratory disease (BRD) with the results of bacterial culture and tulathromycin susceptibility from isolates of deep nasopharyngeal swabs (DNS) as well as viral polymerase chain reaction (PCR) results from nasal swabs revealed poor correlation of bacterial culture and tulathromycin susceptibility with response to tulathromycin metaphylaxis or treatment. 1031 heifers, assumed to be at high-risk (>40% expected BRD morbidity rates), were procured and transported to a research feedlot in Texas. Isolation rates from DNS collected on arrival and at first treatment respectively were: *Mannheimia haemolytica* (10.9% & 34.1%); *Pasteurella multocida* (10.4% & 7.4%); *Mycoplasma bovis* (1.0% & 36.6%); and *Histophilus somni (*0.7% & 6.3%). Prevalence of BRD viral nucleic acid on nasal swabs collected at first treatment were: PI-3V (34.1%); BVDV (26.3%); BoHV-1 (10.8%); and BRSV (54.1%). Increased relative risk of treatment failure was associated with positive viral PCR results, PI-3V (1.2644), BVDV (1.3917), BHV-1 (1.5156), and BRSV (1.3474) from nasal swabs collected at first pull and culture of *M. haemolytica* (1.2284) from DNS collected at arrival; however, no other statistically predictable risk of treatment outcomes were measured from DNS for bacterial isolation or tulathromycin susceptibility for *M. haemolytica* or *P. multocida* at arrival or first treatment. Predictive values of bacterial culture and tulathromycin susceptibility were substantially lower than the 85% level expected with susceptibility testing. These results indicate tulathromycin susceptibility testing of isolates of *M. haemolytica* or *P. multocida* from DNS collected on arrival or at first pull unreliably predict clinical efficacy of tulathromycin for BRD control or treatment most likely due to impacts of unpredictable risk factors and other viral and/or bacterial BRD comorbidities.

## Introduction

For many decades, Bovine Respiratory Disease (BRD) has been a frustrating problem of cattle, veterinarians, and producers due to complex interactions that exist between multiple bacterial and viral pathogens, i.e., polymicrobial infections, inconsistent environmental and management risk factors, and variable immune capabilities of cattle that can alter the outcome of BRD and make diagnosis and treatment frustrating.^1,2,3^ Some consider immune compromise to be as important in the disease process of BRD as the various infectious pathogens involved.^4,5,6^ Together, these interactions as well as limitations of sampling lungs of live cattle, contribute to the lack of a “gold standard” for the definitive diagnosis of BRD. Currently, the most commonly applied method for diagnosing BRD in the cattle industry of North America, i.e. the “industry standard”, is the use of clinical signs, even though variations in ability to observe clinical signs of individual animals in a group along with an inability to observe subclinical signs makes it an imperfect standard.^7^ Bacterial culture and antimicrobial susceptibility testing are commonly used diagnostic test methods to help identify bacterial pathogens involved in BRD which practitioners use as a guide to select or evaluate BRD antimicrobial treatments.^7,8^ However, limitations of these methods specifically for complex disease processes like BRD, can lead to erroneous conclusions, similar to what has been found in humans suffering from polymicrobial disease.^9^

Polymerase chain reaction (PCR) is a diagnostic technique that amplifies targeted regions of nucleic acid for the detection and diagnosis of infectious diseases as well as other applications.^10^ Viral PCR is gaining favor as a diagnostic tool because of the ease of application and recent advances in technology that provide relatively inexpensive and rapid test results. The usefulness of PCR testing has benefits of the ease of application and cost but also drawbacks with interpretation and validation of results for BRD. One of the challenges with diagnosis of BRD is obtaining samples from the lower respiratory tract in live cattle, because it involves invasive, time consuming, and intricate procedures that are not routinely performed or widely practiced in the field and have variable inherent risks to the animal. Consequently, sampling the upper respiratory tract is currently used more frequently for BRD diagnostic testing. The validity of diagnostic tests for BRD from samples taken via the upper respiratory tract has been investigated by many different researchers with inconsistent conclusions.^11–23^ The advantages of PCR testing may help answer some unanswered questions about BRD.

Challenges that BRD presents for these diagnostic test methods, i.e., inconsistent polymicrobial etiologies and issues with collecting lower respiratory tract samples, cause some to believe these methodologies are less useful. The primary objective of this study was to evaluate the agreement between tulathromycin susceptibility test results of *Mannheimia haemolytica (M. haemolytica),* and *Pasteurella multocida (P. multocida),* isolates derived from DNS with clinical outcome in high-risk feeder cattle, when administering tulathromycin for BRD treatment or metaphylaxis. A secondary objective of the study was to assess the agreement with multiplex PCR, including bovine viral diarrhea virus (BVDV), parainfluenza-3 virus (PI-3V), bovine herpes virus-1 (BHV-1), and bovine respiratory syncytial virus results (BRSV), from nasal swabs at first pull to the clinical outcome of high-risk feeder heifers with BRD using clinical scoring (CAS) assessed by an experienced investigating veterinarian.

## Materials and Methods

This prospective cross-sectional cohort study observed eleven truckloads (94 – 116 heifers/load) procured primarily from sale barns in Alabama, but also Kentucky, and South-Central Texas for 42 days following tulathromycin metaphylaxis and treatment for BRD. Agreement of bacterial culture, tulathromycin susceptibility testing, and viral PCR was compared with tulathromycin metaphylaxis and treatment outcomes. Based on previous history of procuring heifers of similar age (6-9 months), weight (205-250 kg.), and origin from these livestock auctions, the goal was to purchase animals at “high-risk” (> 40% morbidity) of developing BRD within 30 days of arrival in the feedlot. This study was executed from March to May at a research feedlot located in the Texas Panhandle with management, facilities and environment like most cattle feeding operations in the Southern & High Plains.

The average body weight of heifers on Day 0 of the study was 226 kg (498 lbs.), ranging between 144 kg (317 lbs.) and 292 kg (642 lbs.). Heifers from each truckload, were allocated between March 5, 2015 and March 28, 2015, to pens of 20 animals with 90 to 120 square feet of pen space and 18 to 24 inches of bunk space. All cattle were housed in dirt floor pens with steel post and cable fencing with concrete fence-line bunks and aprons and float activated water troughs, representative of many North American feed yards. The cattle were fed a total mixed starter ration consisting of 47.5% steam flaked corn, 23% ground alfalfa hay, 5% supplement, 5% molasses, 4.5% cotton seed meal, 15% cotton seed hulls. Feed, delivered with a mixer truck with load cells, was fed twice a day with amounts adjusted daily, so the cattle consumed all the feed by the next morning, i.e., slick bunk management. Within 36 hours of arrival, 1031 high-risk feeder heifers were enrolled in the study (Day-0). Animals with signs of lameness or disease other than BRD which the investigating veterinarian expected would prevent them from finishing the study were excluded before enrollment. The investigating veterinarian walked through each pen and scored each individual animal in each pen at approximately the same time each day for a period of 42 days using a customary clinical appearance score (CAS) system **(Table 1)**. Identification of all animals with a CAS>0 was recorded daily on written forms and observations on the last truckload of cattle ended on May 9, 2015. Written forms were submitted to the statistician for verification and transfer to the software program for data analysis.

**Table 1.** CLINICAL APPEARANCE SCORE INDEX (CAS)

| **CAS Score** | **Appearance Descriptors** |
| --- | --- |
| **0**<br>**No BRD** | No BRD clinical signs; calf is healthy and Bright, Alert, and Responsive when approached. |
| **1**<br>**Mild BRD** | Calf looks ill until approached by observer (calf brightens up, moves readily and appears normal); if observer loses sight of calf and upon re-exam of the pen, he/she CANNOT readily locate and ID calf as ill.<br>Mild depression; slower in movement but no signs of weakness; small amount of serous nasal discharge; slight ocular discharge. |
| **2**<br>**Moderate BRD** | Calf is obviously ill with BRD; the calf shows no change in appearance when approached by observer (does not brighten up and moves slowly or reluctantly). If observer loses sight of calf, upon re-exam of pen, he/she CAN easily locate and ID calf as ill.<br>Moderate depression; signs of weakness or “knuckling” and calf may be reluctant to stand or move about pen; some shallowness apparent in left flank; considerable serous nasal discharge or moderate amount of mucopurulent nasal discharge; dyspnea or respiratory rate is increased; cough or coughing episodes are present. |
| **3**<br>**Severe BRD** | Calf is severely ill with BRD; the calf shows no change in appearance when approached by observer; abnormal respiration and/or depression present.<br>Severe depression; stumbling or moves with extreme prodding; obvious lack of fill in left flank signaling anorexia; head lowered or extended to facilitate breathing; may be open-mouthed breathing with considerable noise (expiratory grunts, moans); copious mucopurulent to purulent nasal discharge; cough or coughing episodes are present.<br>Administer emergency intervention therapy. |
| **4**<br>**(Moribund)** | Calf is moribund and near death – calf in general is not ambulatory, cannot rise from recumbency, and cannot acquire food or water. Very likely cannot be removed from pen for treatment without mechanical transport.<br>Euthanize humanely. |

Power calculation was based on the expected incidence of tulathromycin resistant *M. haemolytica* isolates to show 85% agreement with treatment failures. The calculation was based on 1000 head sampled at arrival and a 30% incidence of *M. haemolytica* isolation and 1% incidence of tulathromycin resistance (3 head) and 275 head sampled at first pull with an incidence of 50% *M. haemolytica* isolation and 30% incidence of tulathromycin resistance (21 head) for a total of 24 animals with tulathromycin resistant *M. haemolytica* isolates. Due to low or no tulathromycin resistance reported in the literature, it was not expected to have enough tulathromycin resistant isolates from arrival samples and therefore the protocol was designed to combine all samples for susceptibility analysis.

Randomization of individual cattle or lots of cattle was not necessary because of the study design, i.e., all cattle received the same arrival protocol **(Table 2)** administered by individual body weight and label instructions. Certified scales were calibrated each day before enrollment of cattle. Laboratory personnel were masked to the origin of samples and research personnel were masked to the laboratory results until the completion of the study. This study complied with all applicable animal welfare regulations related to the humane care and use of animals and all procedures and products were reviewed and approved by an animal use and welfare committee. Any animal identified with critically severe clinical signs (CAS 3), was given immediate emergency therapy and removed from the study. Any animal found to be moribund (CAS 4), was removed from the study and humanely euthanized per the American Veterinary Medical Association “Guidelines for Euthanasia of Animals,” 2013 Edition.^24^ Products in the arrival and treatment protocols were administered per Beef Quality Assurance Guidelines (BQA). **(Table 2)** A timeline of the study design is illustrated in **Figure 1.**

**Table 2.**
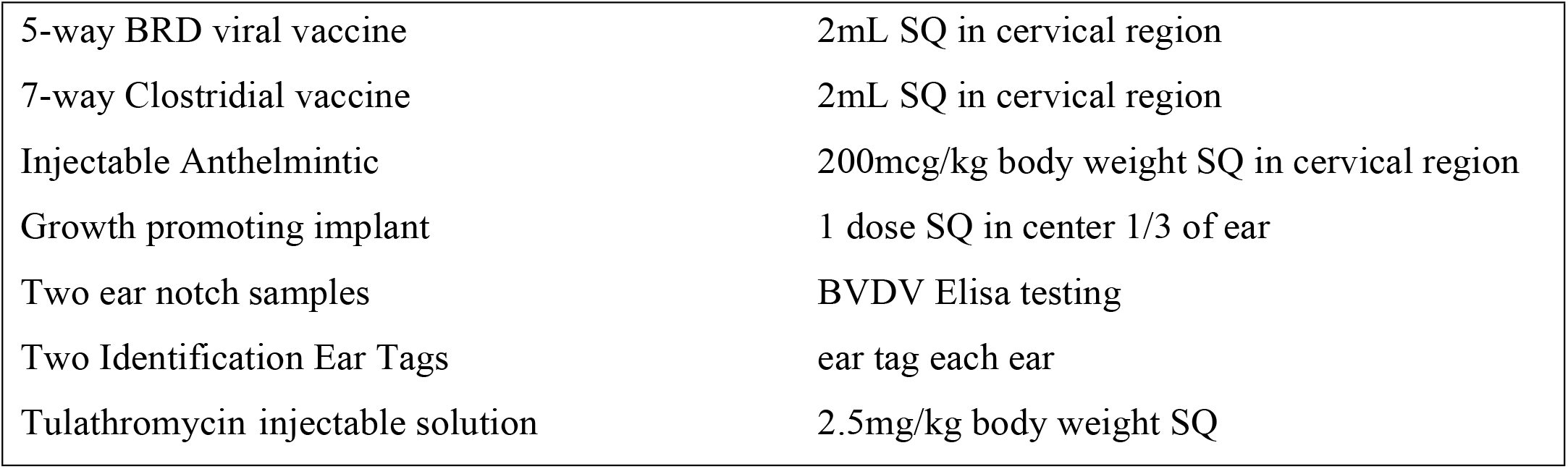
ARRIVAL PRODUCTS AND PROCEDURES

**Figure 1.**
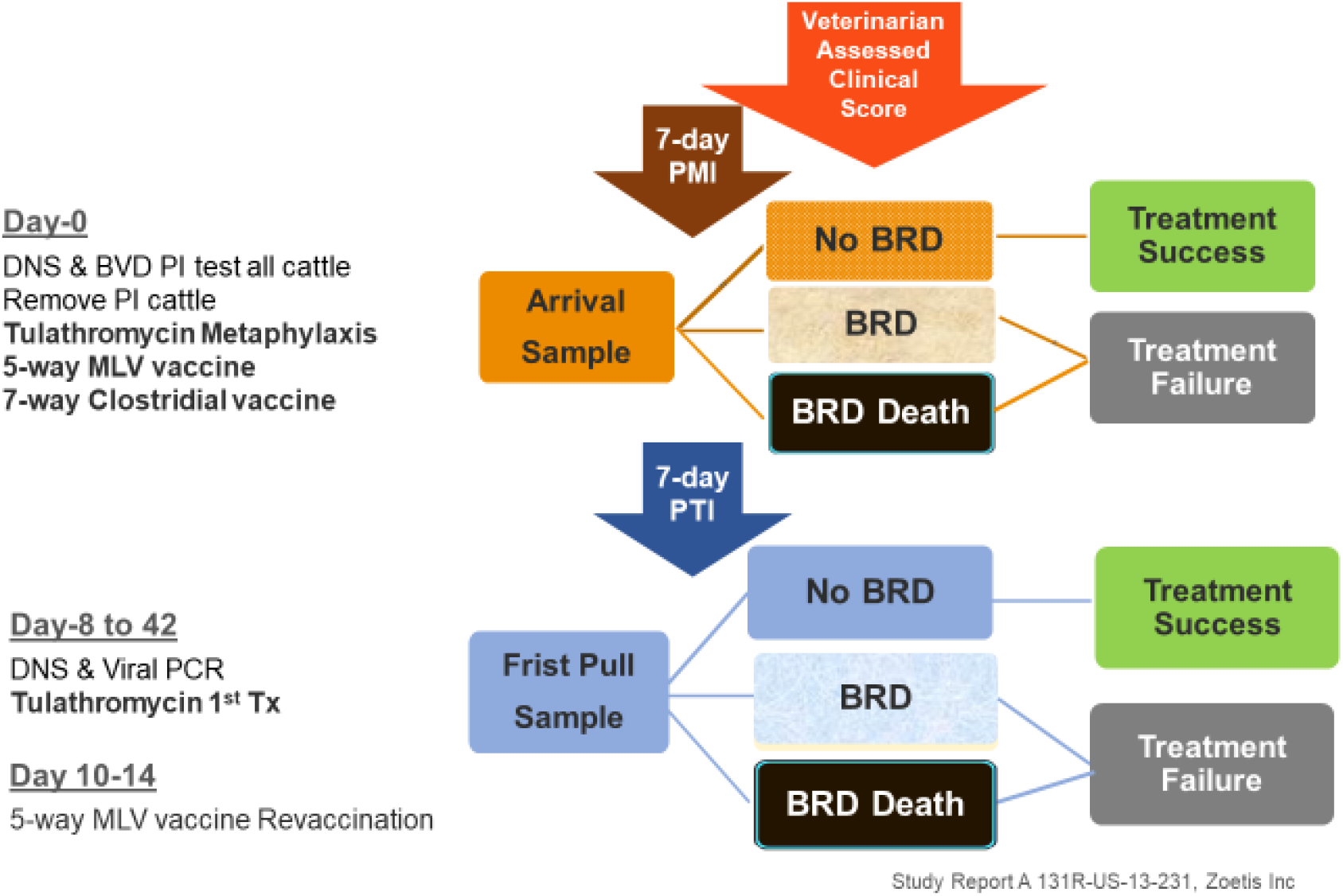
STUDY TIMELINE

DNS specimens were shipped to an accredited research laboratory in Amies transport medium containing charcoal and packed with ice packs via overnight courier, except samples collected on Saturday and Sunday which were held at 2-8oC and shipped on Monday. Samples were process on the date of receipt in the lab, except samples received on Saturday which were held at 2-8oC until the following business day. All samples were received at the lab, cooled, in good condition and were labeled with the study number, date of collection, sample type, and animal ID. DNS samples were cultured for identification of *Mannheimia haemolytica, Pasteurella multocida,* and *Histophilus somni (H. somni).* Specimens were streaked for isolation on a 5% sheep blood agar plate (BA) and to modified Hayflick’s agar (HFA). The BA plates were incubated overnight, and the HFA plates were incubated up to 7 days at 36±2oC in 5±2% CO_2_. The incubated BA plates were observed for the presence of presumptive *H. somni, M. haemolytica*, and *P. multocida* colonies. The HFA agar plates were observed for typical *Mycoplasma bovis (M. bovis)* colonies. Each colony with a presumptive identity of *H. somni, P. multocida*, or *M. haemolytica* was identified by Maldi Biotyper. Presumptive *M. bovis* colonies were purified by two serial passages on HFA. Presumptive colonies were then dienes stained and tested for inhibition by digitonin to identify the isolates as *Mycoplasma* species. Speciation of *Mycoplasma* isolates was performed using a validated PCR procedure to confirm an identification of *Mycoplasma bovis*.

Tulathromycin susceptibility, using MIC broth microdilution technique, was determined on *M. haemolytica* and *P. multocida* isolates using one representative colony from each sample. Tulathromycin minimum inhibitory concentrations (MIC) values for *M. haemolytica* and *P. multocida* were determined using plates prepared by the research laboratory on May 29, 2015 which contained doubling dilution concentrations of tulathromycin from 0.12-64 ug/mL. Positive and negative growth control wells were included for each dilution series. MIC tests were performed using Clinical & Laboratory Standards Institute (CLSI) procedures.^25^ Swabs were streaked onto 5% sheep blood agar and modified Hayflick agar and incubated in 5 ±2% CO_2_ at 36 ±2oC overnight for the blood agar plates and up to 7 days for the Hayflick agar plates. Incubated plates were observed for typical colonies of *H. somni, M. haemolytica, P. multocida,* and *M. bovis*. Cation adjusted Mueller Hinton broth (MHB) was used for *P. multocida* and *M. haemolytica* isolates. Plates were incubated aerobically at 36±2oC for 19.5 hours. The MGB quality control organisms, *Enterococcus faecalis* and *Staphylococcus aureus*, tested on each testing date, were incubated aerobically at 36±2oC for 19.0-19.5 hours. Only one isolate was tested from each sample unless presumptive identification of the isolate was not confirmed.

Animals were not eligible for first pull treatment until the eighth day (7-day PMI) after metaphylatic/arrival tulathromycin administration. Starting on day eight, animals with a CAS=1 and a rectal temperature >39.7o C or a CAS ≥2 (regardless of rectal temperature) were eligible to receive first pull treatment for BRD and were classified as treatment failures from the metaphylaxis tulathromycin administration. At first pull treatment, a second DNS for culture and tulathromycin sensitivity along with a nasal swab for multiplex viral PCR was collected and tulathromycin was administered. First pull procedures, as listed in **Figure 1,** included:

- DNS collected using the same method as day-0.
- A nasal swab^i^ collected from the external nares after cleaning the nares with a clean paper towel. Following collection, nasal swabs were stored in red stopper tubes^ii^, refrigerated, and shipped with ice packs, overnight to an accredited laboratory for multiplex PCR testing.
- Tulathromycin administered for the treatment of BRD per body weight and label instructions following BQA guidelines.

Upon arrival at the laboratory, dry nasal swabs were suspended in minimal essential medium and then frozen until testing was performed as described. Nucleic acid was purified from the submitted swab sample using the MagMAX-96 Viral RNA isolation kit from ThermoFisher. Briefly, the swab was moistened in ∼700uL 1X phosphate buffered saline, pH 8.0 and ∼150uL utilized for nucleic acid extraction. Sample was combined with 20uL of magnetic bead mix 10uL lysis binding enhancer and 10uL RNA binding beads and 400uL lysis binding solution 200uL lysis binding concentrate, 1uL carrier RNA (1ug/ul), 1uL XIPC RNA (at 10,000 copies/uL), and 200uL 100% isopropanol in a 96-well deep-well plate which was labeled as the sample plate. Nucleic acid extraction was performed using a KingFisher 96 automated particular processor. The following plates were added to the KingFisher 96 for extraction: sample plate, wash solution 1 plate (300uL/well), wash solution 2 plate (300uL/well) and elution buffer plate (90uL/well). Following nucleic acid extraction, eluted nucleic acid was kept refrigerated prior to PCR setup.

BVD, BRSV and BPI3 were screened via multiplex PCR, utilizing the ThermoFisher PathID Multiplex One Step RT-PCR kit according to the manufacturer’s instructions, along with primers and probes for detection of BVD, BRSV, BPI3 and XIPC (an exogenous internal control). Each reaction contained the following: 12.5uL 2X Multiplex RT-PCR buffer, 2.5uL 10X multiplex enzyme mix, 1uL 25X primer-probe mix (containing all oligonucleotides for detection of all four targets) and 1uL nuclease free water; 8uL of extracted nucleic acid was added to 17uL of MasterMix for a total 25uL reaction volume. The RT-qPCR was performed using the Applied Biosystems 7500Fast instrument. Cycling parameters were as follows: 48°C for 10min (1 cycle), 95°C for 10min (1 cycle), and 40 cycles at 95°C for 15sec and 55°C for 45sec. Samples with a quantification cycle (Cq) ≤ 37.0 were considered positive for the above-mentioned targets.

Detection of IBR was assessed in a separate PCR using the PathID qPCR MasterMix from Thermo Fisher, along with primers and probes for the detection of IBR and XIPC. Each reaction contained the following: 12.5uL 2X PathID qPCR buffer, 1uL 25X primer-probe mix (containing the oligonucleotides for the detection of IBR and XIPC) and 3.5uL nuclease free water; 8uL of extracted nucleic acid was added to 17uL of MasterMix for a total 25uL reaction volume. Cycling parameters and quantification cycle cutoff were the same as for the multiplex assay above.

Animals were not eligible for additional treatment of BRD for a period of 7 days after first pull administration of tulathromycin (7-day PTI) unless, per protocol, animals that scored a CAS≥3 were given emergency treatment or were euthanized. All animals that were euthanized or died during the study were necropsied by the investigating veterinarian and further testing, e.g., histopathology, IHC, culture, PCR, viral isolation, was performed, if necessary, to determine a definitive diagnosis for death.

Data was transferred from written forms, validated, stored, and analyzed in a centralized data management system, SAS version 9.3. Data was summarized in contingency tables and analyzed for *Sensitivity, Specificity, Positive Predictive Value* (PPV), *Negative Predictive Value* (NPV), and *Relative Risk of Treatment Failure (RRTF)*. Clinical outcome was observed for 42 days on each load of cattle and defined as ***Treatment Success***if the animal did not have clinical signs of BRD (CAS-0) during the 42-day observation period, according to the CAS assessed daily by the investigating veterinarian. ***Treatment Failure***was defined as an animal with clinical signs of BRD (CAS≥1)assessed by the investigating veterinarian using the CAS any time after the 7-day treatment suspension period or death due to BRD. Animals classified as treatment failures were given additional antibiotic treatment according to the protocol or euthanized (CAS≥3).

For analysis of sensitivity, specificity, positive and negative predictive value and relative risk of treatment failure for bacterial culture and multiplex PCR testing **Table 3**, *True Positive* result was defined as animals with identification of BRD bacterial and/or viral pathogens that were classified as treatment failures (died or needed further treatment per CAS) following the 7-day treatment suspension period. *True Negative* result for the same statistical tests was defined as animals with no identification of BRD pathogens that were classified as treatment successes (needing no further treatment per CAS) during the 42-day study. *False Positive* results were defined as animals with identification of BRD bacterial and/or viral pathogens that were subsequently classified as treatment successes and *False Negative* results were defined as animals without evidence of BRD pathogens but classified as treatment failures following the 7-day treatment suspension period.

**Table 3.**
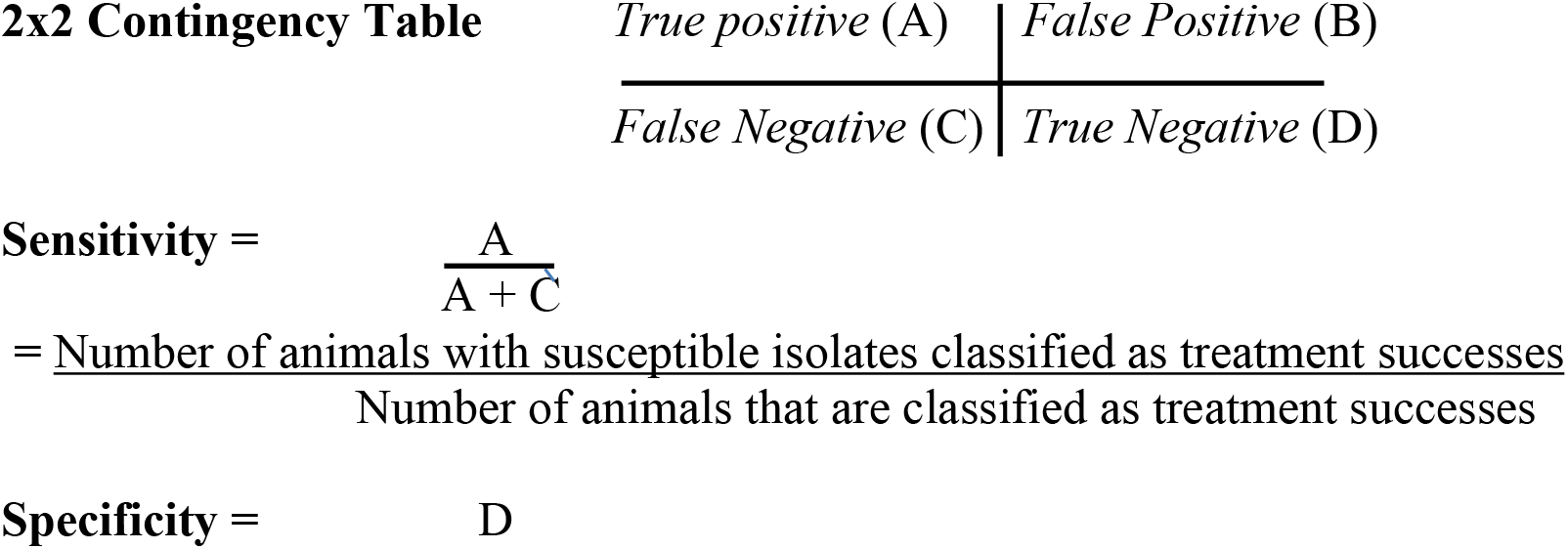
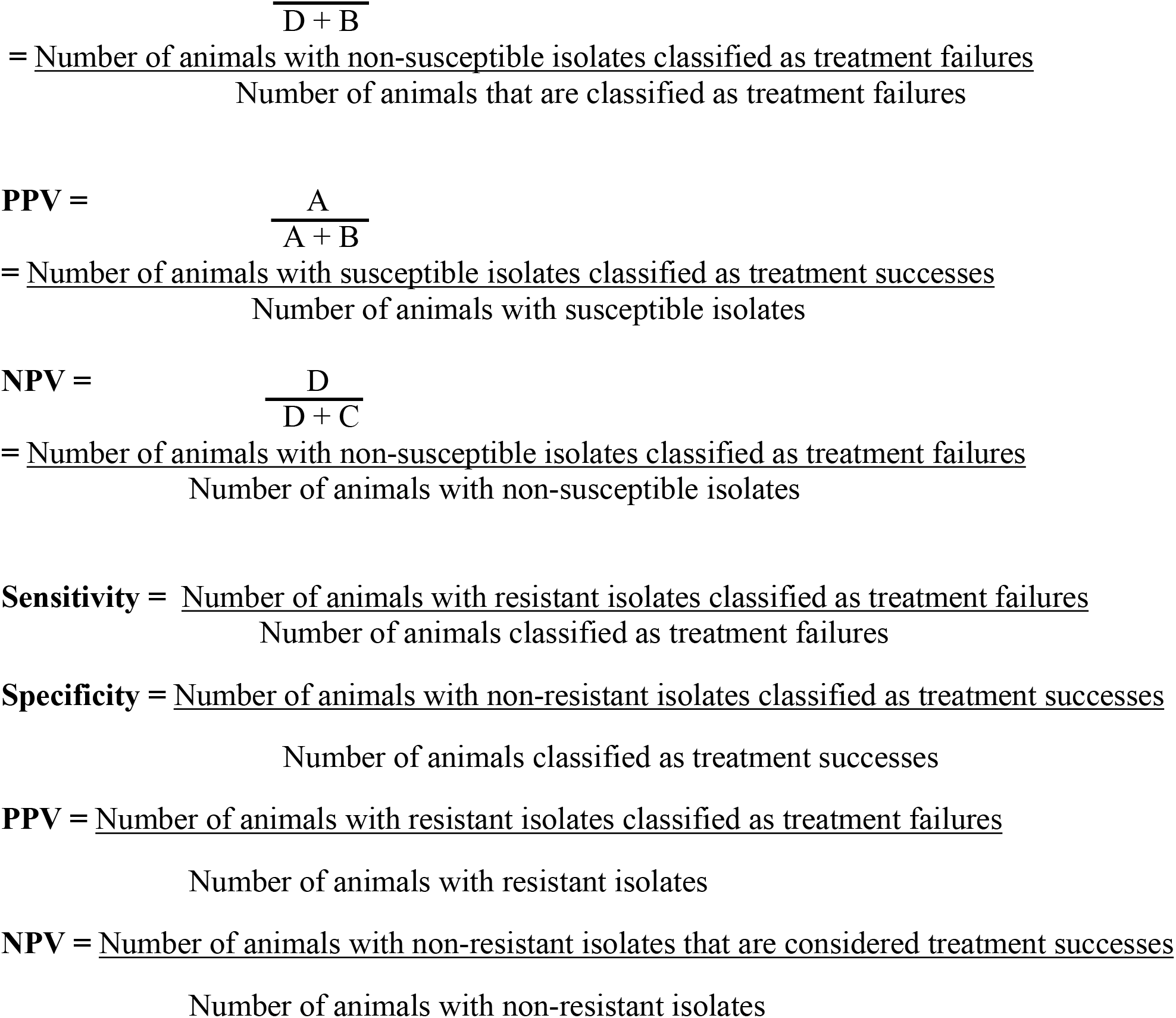
ANALYSIS EQUATIONS

For analysis of sensitivity, specificity, positive, and negative predictive value of tulathromycin susceptibility, CLSI established tulathromycin veterinary specific interpretive criteria for *M. haemolytica* and *P. multocida* were used to establish susceptible (≤ 16 ug/ml), intermediate (32 ug/ml), and resistant (≥ 64 ug/ml) isolates. *True Positive* results were defined as cattle with non-resistant (MIC < 64ug/ml) *M. haemolytica* or *P. multocida* isolates that were classified as treatment successes following no subsequent identification of clinical signs of BRD by the investigating veterinarian during the 42-day study. *True Negative* tulathromycin susceptibility results were defined as any cattle with resistant (MIC ≥ 64ug/ml) *M. haemolytica* or *P. multocida* isolates classified as treatment failures after reoccurrence of clinical signs of BRD following the 7-day treatment suspension period or death due to BRD subsequent to tulathromycin treatment. *False Positive* tulathromycin susceptibility results were defined as all cattle with non-resistant (MIC < 64ug/ml) *M. haemolytica* or *P. multocida* isolates that were classified as treatment failures after identification of clinical signs of BRD by the investigating veterinarian following the 7-day treatment suspension interval. *False Negative* tulathromycin susceptibility results were defined as any cattle with resistant (MIC ≥ 64ug/ml) *M. haemolytica* or *P. multocida* isolates that were classified as treatment successes with no identification of subsequent clinical signs of BRD by the investigating veterinarian during the 42-day study.

## Results

1031 head (eleven truckloads) of English, Continental, and crossbred feeder heifers presumed to be at high risk of developing BRD (>40% morbidity) were enrolled on Day-0 of the study. Mean arrival weight of animals with treatment failure was 225 kg. (495 lbs.), with standard deviation of 22 kg. (49 lbs.) and mean arrival weight of animals with treatment success was 227 kg. (499 lbs.), with standard deviation of 25 kg. (56 lbs.). Mean day-0 weight of first pull cattle with treatment failure was 224 kg. (493 lbs.), with standard deviation of 21 kg. (46 lbs.) and mean arrival weight of animals with treatment success was 227 kg. (499 lbs.), with a standard deviation of 24 kg. (52 lbs.).

Three heifers were found to be persistently infected with BVDV, with agreement of both labs, and were removed from the study within the first 5 days. After the initial 7-day treatment suspension period (PMI) following the metaphylaxis administration of tulathromycin, 402 heifers (38.9%) were identified by the investigating veterinarian as having clinical signs of BRD, thus classified as treatment failures of tulathromycin metaphylaxis. Mean day of first pull was 13 days with a range per load of 8-16 days. Morbidity per load ranged from 23% to 44% and treatment failure ranged from 23% to 49% after tulathromycin metaphylaxis and 38% to 69% after first pull tulathromycin therapy. There were three instances where animals were given BRD treatments that deviated from the protocol resulting in removal of two animals from the study, after initial processing, and the removal of three animals after first pull BRD treatment which resulted in 1026 total animals in the arrival population and 399 animals in the first pull population. No animals were treated on an emergency basis or euthanized because of debilitating clinical signs of BRD. Isolation rates for isolates of *M. haemolytica, P. multocida, H. somni,* and *M. bovis* collected from DNS at arrival and first treatment are summarized in **Figure 2.** Prevalence of *M. haemolytica* (10.9% and 34%), *M. bovis* (1.0% and 36.6%) and *H. somni* (0.6% and 6.3%) all substantially increased in cattle sampled at first pull compared to the arrival sampling however, prevalence of *P. multocida,* decreased slightly when cattle were sampled at first pull treatment (7.4%) compared to cattle cultured at arrival (10.4%).

**Figure 2.**
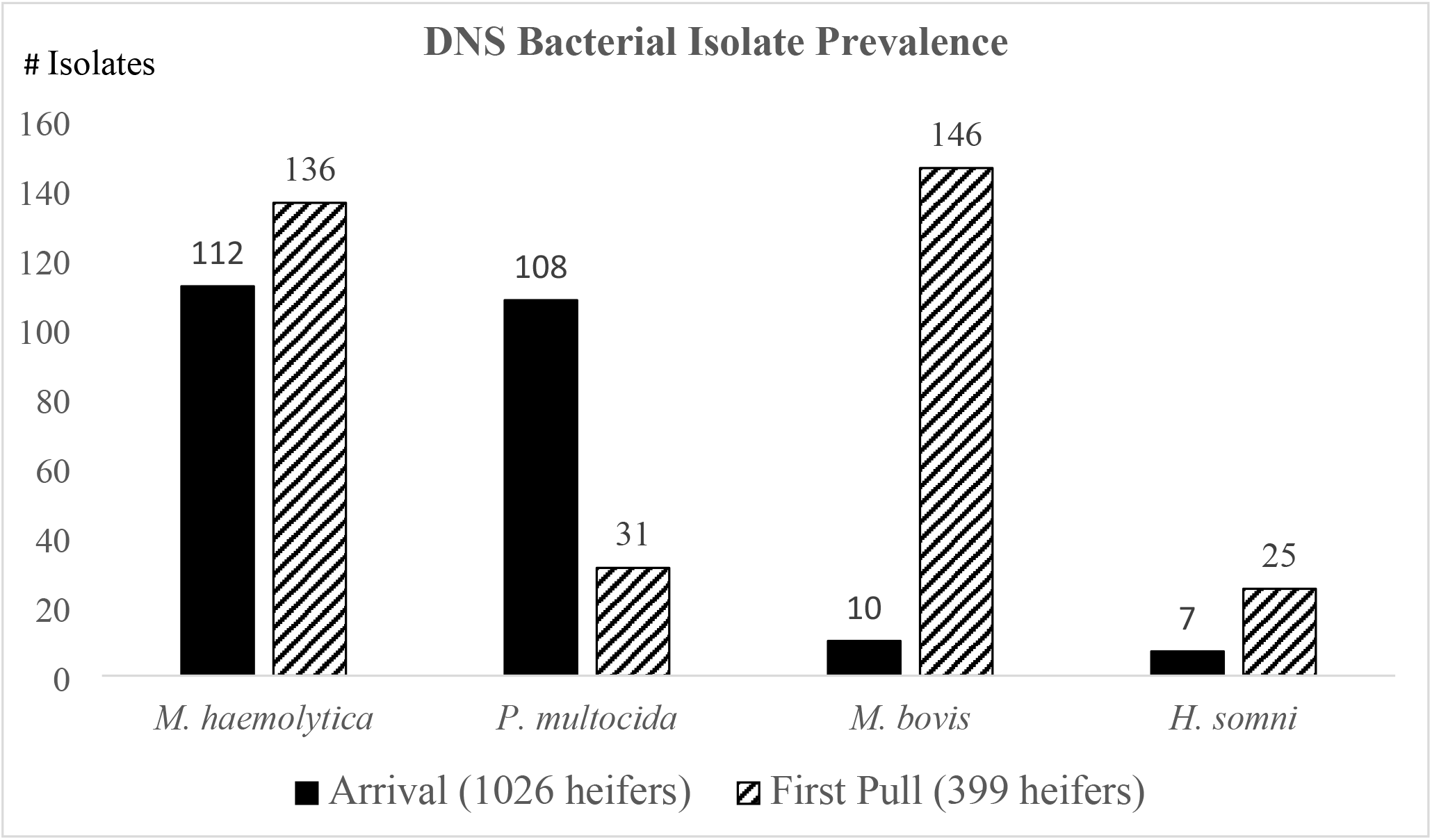
PREVALENCE OF BRD PATHOGENS ISOLATED FROM DNS IN HIGH-RISK FEEDER HEIFERS AT ARRIVAL AND FIRST PULL

Frequency distributions of MICs for all *M. haemolytica* and *P. multocida* isolates collected via DNS are summarized in **Figure 3**. *M. haemolytica* isolates tested were bimodally distributed between isolates that were susceptible or resistant to tulathromycin. There was a low prevalence of *M. haemolytica* isolates intermediately resistant (0.7%) to tulathromycin and *P. multocida* isolates were 85% susceptible to tulathromycin.

**Figure 3.**
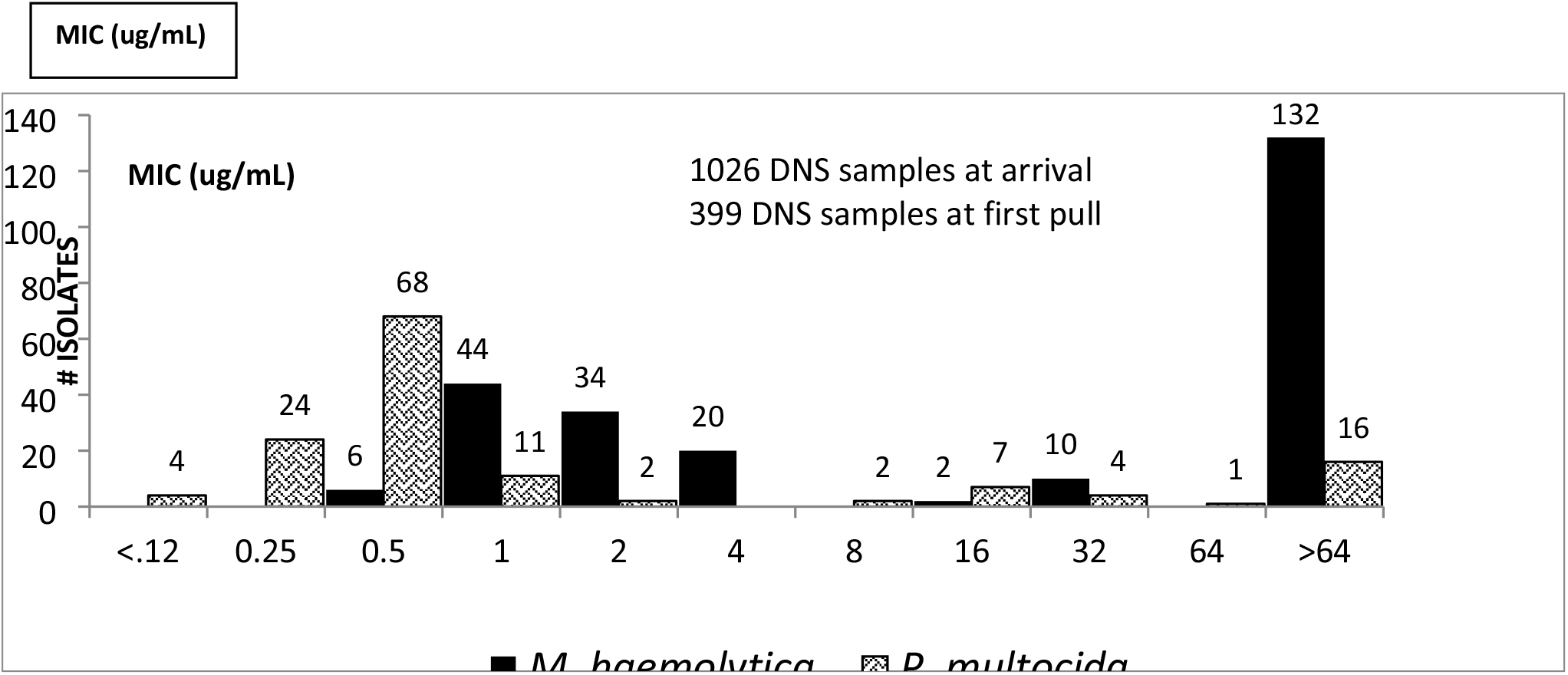
MIC FREQUENCY DISTRIBUTION OF *M. HAEMOLYTICA* AND *P. MULTOCIDA* ISOLATES FROM DNS SAMPLES OF HIGH-RISK FEEDER HEIFERS AT ARRIVAL AND FIRST PULL

Treatment Success Rates (TSR) for cattle with or without *M. haemolytica* or *P. multocida* isolated via DNS are summarized in **Figure 4.** TSRs were similar except for cattle with isolates of *M. haemolytica* (112/1026 head) from DNS collected at arrival which had 9% lower TSR than cattle without isolation of *M. haemolytica* (914/1026 head). Lower TSR for cattle with isolation of *M. haemolytica* on arrival was associated with an increased risk of treatment failure (p value 0.049). Cattle with isolates of *M. haemolytica* (134/399 head), without isolates of *M. haemolytica* (265/399 head), with *P. multocida* (29/399), without *P. multocida* (370/399), with *H. somni* (25/399), without *H. somni* (374/399), with *M. bovis* (79/399) or without *M. bovis* (320/399), cultured from DNS collected at first pull all had similar TSR. **Figure 4**.

**Figure 4.**
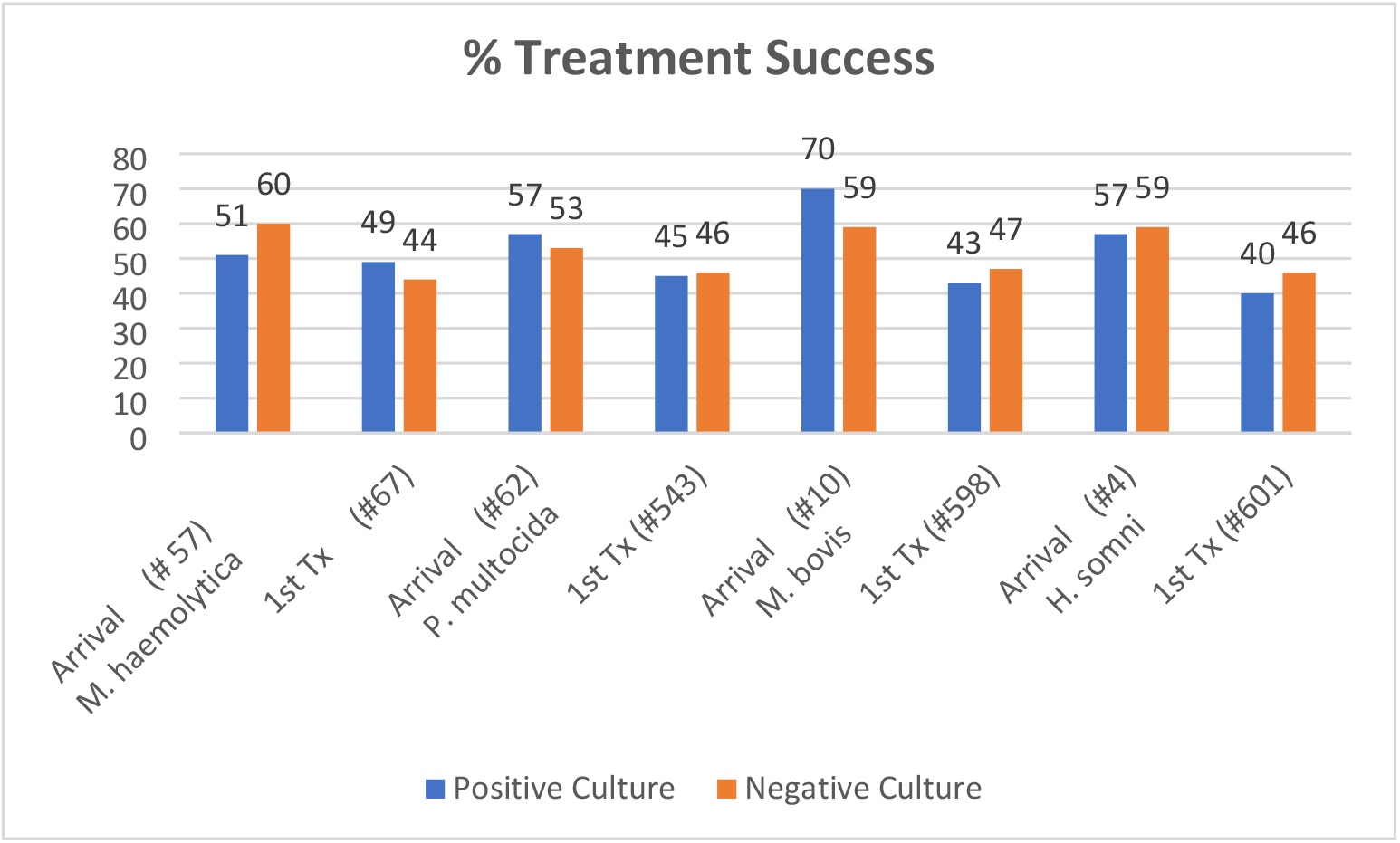
HIGH-RISK FEEDER HEIFER TREATMENT SUCCESS PROPORTION BY CULTURE STATUS AND TIMING OF SAMPLE

**Figure 5.** illustrates TSRs for cattle with *M. haemolytica* isolates were similar regardless of their tulathromycin susceptibility. Contrary to susceptibility parameters predicting ≥85% of resistant isolates will result in treatment failure and ≥85% of susceptible isolates will result in treatment success, cattle in this study, with *M. haemolytica* isolates susceptible to tulathromycin, had a greater Treatment Failure Rate (TFR) (52%) than cattle with tulathromycin resistant *M. haemolytica,* (50% TFR). MICs for *P. multocida* isolates were associated with lower TFR when comparing susceptible (S) isolates (42.7% TFR) to non-susceptible (I & R) isolates (54.5% TFR) and resistant (R) isolates (50% TFR) compared to non-resistant (S,I) isolates (43.8% TFR) but were far from the ≥85% parameters of CLSI breakpoint predictions.

**Figure 5.**
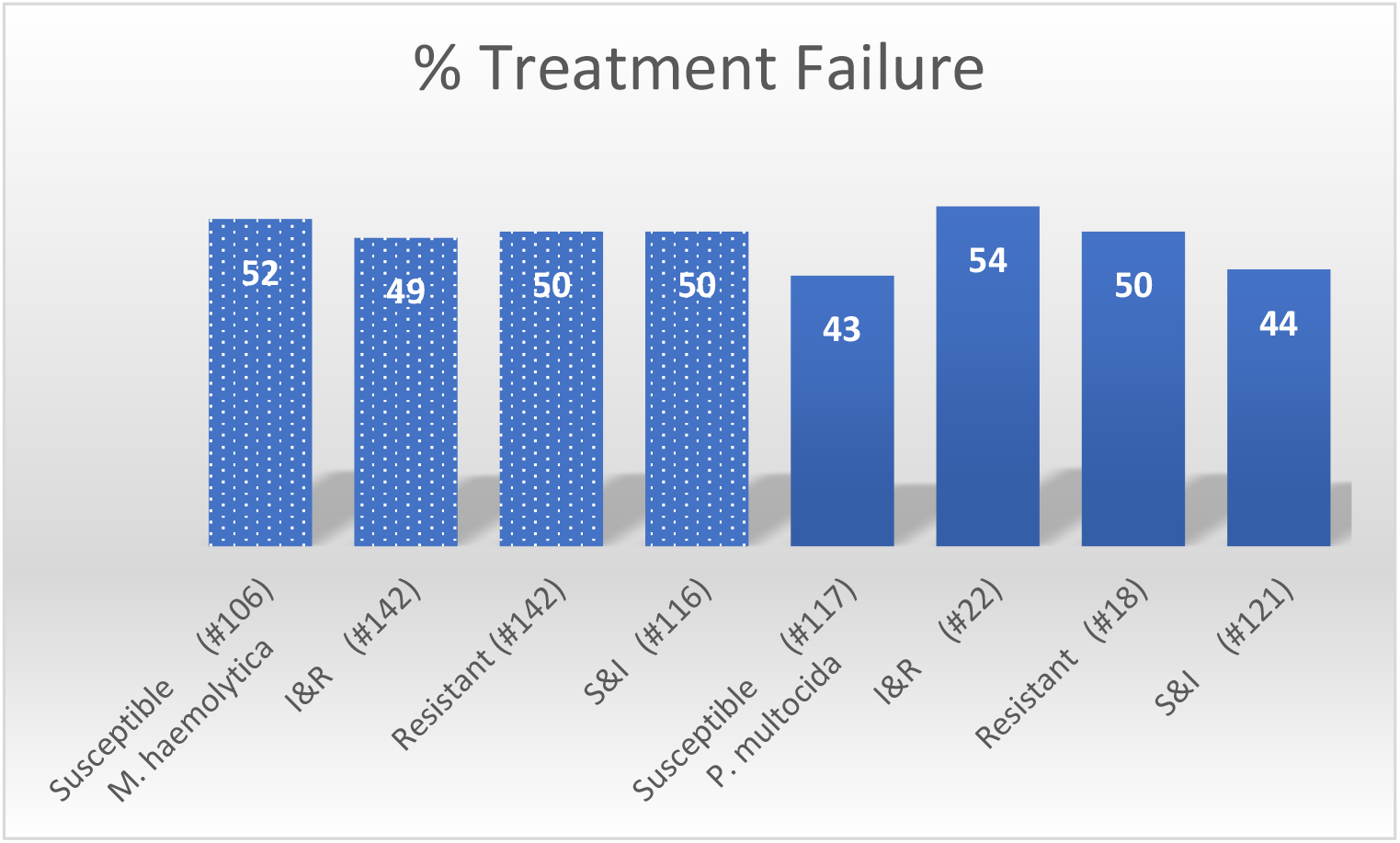
HIGH-RISK FEEDER HEIFER BRD TREATMENT FAILURE BY BACTERIAL ISOLATE AND TULATHRYMYCIN SUSCEPTIBILITY

Prevalence of both susceptible (223/387) and resistant (150/387) isolates was enough to provide meaningful analysis of the data. **Table 4.** The positive and negative predictive values for both susceptible or resistant isolates of either *P. multocida* or *M. haemolytica* were all close to 50%. Comparing **resistant isolates (R) vs. non-resistant (S, I)** isolates of *P. multocida* or *M. haemolytica,* indicated poor sensitivity (.529) and specificity (.468) for *M. haemolytica* isolates and low sensitivity (.145) but high specificity (.883) for *P. multocida* isolates. Comparing **susceptible isolates (S) vs. non-susceptible (I , R)** of the same bacterial pathogens revealed poor sensitivity (.411) and specificity (.556) for *M. haemolytica* isolates and good sensitivity (.870) but poor specificity (.193) for *P. multocida* isolates, due to greater number of intermediate isolates of *P. multocida*.

**Table 4.**
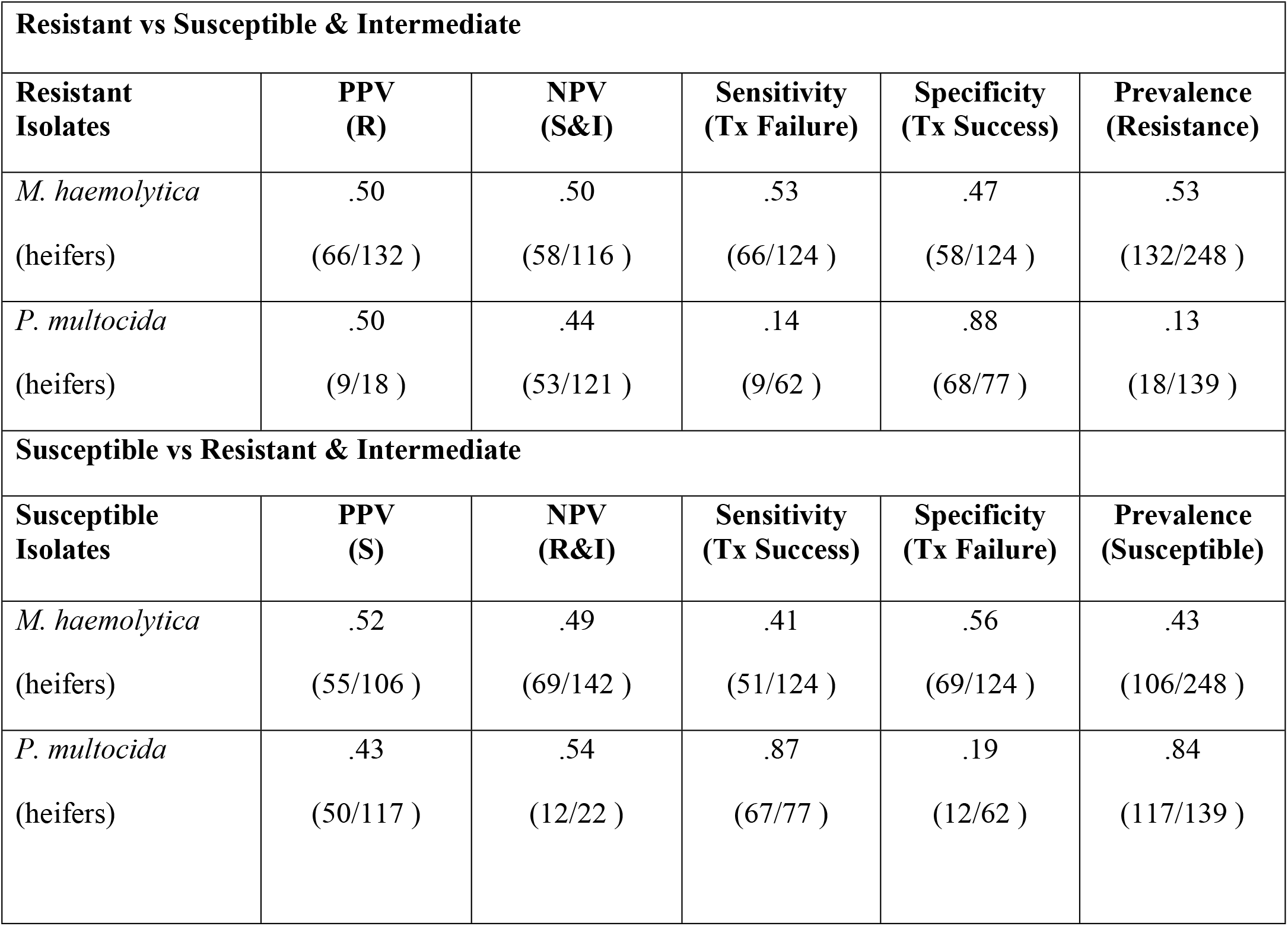
SENSITIVITY, SPECIFICITY, PREDICTIVE VALUE, AND PREVALENCE OF TULATHRYMYCIN RESISTANCE FOR ALL *M. HAEMOLYTICA* AND *P. MULTOCIDA* ISOLATES COLLECTED FROM DNS SAMPLES FROM HIGH-RISK FEEDER HEIFERS

Frequency distribution of multiplex PCR results from nasal swabs collected at first pull treatment is summarized in **Table 5.** Prevalence of positive PCR results for the 4 common respiratory viruses was 54% for BRSV, 34% for PI-3V, 27% for BVDV, and 11% for BoHV-1, in this study population of high-risk feeder cattle. **Table 6,** shows that high-risk feeder heifers with positive viral PCR results from nasal swabs, had a higher percentage of treatment failure (ranging from 61% to 77%) compared to cattle that were negative to viral PCR at first pull (45 to 51%). **Table 9**, indicates that the high-risk feeder heifers with positive viral PCR results from nasal swab samples taken at first pull had significantly greater risk of treatment failure (p-values .01-.000l). Data in **Table 8,** suggests acceptable PPV for BHV-l (.77), BVDV (68), PI3V (.62), and BRSV (.61) and good specificity for BHV-1 (.95), BVDV (.81), and PI3V (.72). However, in this study, negative viral PCR results showed poor agreement with treatment success validated by poor NPV (.49-.55). Data in **Table 7,** implies, as the number of pathogens isolated per heifer increased, the proportion of cattle with treatment failure also increased. Greater frequency of one to three pathogens per animal was measured in this population with a median of two pathogens. Less than twenty-five percent of the cattle showing signs of BRD at first pull, had only one BRD pathogen isolated which emphasizes the polymicrobial involvement of BRD.

**Table 5.** FREQUENCY OF VIRAL MULTIPLEX PCR FROM NASAL SECRETIONS COLLECTED AT FIRST PULL TREATMENT

| (399 heifers tested) | Negative<br>PCR | % | Positive<br>PCR | % |
| --- | --- | --- | --- | --- |
| **Parainfluenza Virus-3** | 263 | 66 | 136 | 34 |
| **Bovine Respiratory Syncytial Virus** | 183 | 46 | 216 | 54 |
| **Bovine Viral Diarrhea Virus** | 294 | 74 | 105 | 27 |
| **Bovine Herpes Virus-1** | 356 | 89 | 43 | 11 |

**Table 6.** PROPORTION OF HIGH-RISK FEEDER HEIFERS WITH TREATMENT FAILURE BY VIRAL PCR

| **Viral Pathogen**<br>**(399 head)** | **Treatment Failure with**<br>**Negative Viral PCR** | **Treatment Failure with**<br>**Positive Viral PCR** |
| --- | --- | --- |
| **PI-3V** (heifers) | 49.4% (130 ) | 62.5% (85 ) |
| **BRSV** (heifers) | 45.4% (83 ) | 61.1% (132 ) |
| **BVDV** (heifers) | 48.8% (143 ) | 67.9% (72 ) |
| **BoHV-1** (heifers) | 34.0% (181 ) | 77.3% (34 ) |

**Table 7.** NUMBER OF PATHOGENS PER HIGH-RISK FEEDER HEIFER ISOLATED AT FIRST PULL WITH BRD TULATHROMYCIN TREATMENT OUTCOME

| **#**<br>**Pathogens** | **Total** | **%** | **#**<br>**Failures** | **%**<br>**Failures** | **#**<br>**Success** | **%**<br>**Success** |
| --- | --- | --- | --- | --- | --- | --- |
| 0 | 45 | 11% | 18 | 40% | 27 | 60% |
| 1 | 87 | 22% | 42 | 48% | 45 | 52% |
| 2 | 125 | 32% | 67 | 54% | 58 | 46% |
| 3 | 81 | 21% | 50 | 62% | 31 | 38% |
| 4 | 42 | 11% | 28 | 67% | 14 | 33% |
| 5 | 8 | 2% | 8 | 100% | 0 | 0% |
| 6 | 6 | 2% | 5 | 83% | 1 | 17% |
| Total | 394 | 100% | 218 | 55% | 176 | 45% |

**Table 8.** SENSITIVITY, SPECIFICITY, PPV, NPV, AND PREVALENCE OF VIRAL PCR FROM NASAL SWAB OF HIGH-RISK HEIFERS AT FIRST PULL

| Microorganism | Sensitivity | Specificity | PPV | NPV | Prevalence |
| --- | --- | --- | --- | --- | --- |
| Bovine Herpes Virus-1<br>(heifers) | .16<br>(34/215 ) | .95<br>(174/184 ) | .77<br>(34/44 ) | .49<br>(174/355 ) | .11<br>(44/399 ) |
| Bovine Viral Diarrhea<br>Virus<br>(heifers) | .33<br>(72/215 ) | .81<br>(150/184 ) | .68<br>(72/106 ) | .51<br>(150/293 ) | .26<br>(106/399 ) |
| Bovine Respiratory<br>Syncytial Virus<br>(heifers) | .61<br>(132/215 hd.) | .54<br>(100/184 ) | .61<br>(132/216 ) | .55<br>(100/183 ) | .54<br>(216/399 ) |
| Parainfluenza Virus-3<br>(heifers) | .39<br>(85/215 ) | .72<br>(133/184 ) | .62<br>(85/136 ) | .51<br>(133/263 ) | .34<br>(136/399 ) |

**Table 9.** RELATIVE RISK OF TREATMENT FAILURE FOR BRD DIAGNOSTIC TEST METHODS FROM UPPER RESPIRATORY TRACT SAMPLES IN HIGH-RISK FEEDEER HEIFERS

| **Bacterial Isolates at Arrival** | **Relative Risk of Treatment Failure** | **95% Confidence Interval** | **p-value** |
| --- | --- | --- | --- |
| *M. haemolytica* | 1.23 | 1.00 – 1.51 | .05 |
| *P. multocida* | 1.03 | .81– 1.30 | .81 |
| *H. somni* | .81 | .26 – 2.52 | .72 |
| *M. bovis* | .73 | .28 – 1.89 | .52 |
| **Bacterial Isolates at First Pull** |  |  |  |
| *M. haemolytica* | .89 | .73 – 1.09 | .25 |
| *P. multocida* | 1.01 | .72 – 1.40 | .97 |
| *H. somni* | 1.01 | .79 – 1.54 | .57 |
| *M. bovis* | 1.08 | .90 – 1.30 | .39 |
| **Resistant Isolates at Arrival** |  |  |  |
| *M. haemolytica* | 1.02 | .25 – 4.13 | .98 |
| *P. multocida* | N/A | N/A | N/A |
| **Resistant Isolates at First Pull** |  |  |  |
| *M. haemolytica* | .75 | .42 – 1.35 | .34 |
| *P. multocida* | 1.12 | .57 - 2.16 | .72 |
| **All Resistant Isolates** |  |  |  |
| *M. haemolytica* | 1.0 | .78 – 1.28 | 1.00 |
| *P. multocida* | 1.08 | .65– 1.81 | .77 |
| **Susceptible Isolates at Arrival** |  |  |  |
| *M. haemolytica* | 1.51 | .86 – 3.89 | .39 |
| *P. multocida* | N/A | N/A | N/A |
| **Susceptible Isolates at First Pull** |  |  |  |
| *M. haemolytica* | N/A | N/A | N/A |
| *P. multocida* | .69 | .33 – 1.48 | .35 |
| **All Susceptible Isolates** |  |  |  |
| *M. haemolytica* | 1.07 | .83 – 1.31 | .61 |
| *P. multocida* | .78 | .51 – 1.21 | .27 |
| **Viral PCR at First Pull** |  |  |  |
| BVDV | 1.39 | 1.17– 1.66 | .0002 |
| PI3V | 1.26 | 1.06 – 1.51 | .01 |
| BHV-1 | 1.52 | 1.25 – 1.83 | .0001 |
| BRSV | 1.33 | 1.11 – 1.63 | .002 |

**Table 9,** summarizes the relative risk of treatment failure (RRTF) with 95% confidence intervals and p-values associated with a positive result for each of the diagnostic procedures analyzed in this study. A value less than 1.0 indicates less relative risk for treatment failure (greater probability of treatment success) and a value greater than 1.0 indicates greater relative risk for treatment failure. 95% confidence intervals with lower and upper limits, less than and greater than 1.0, indicate that the relative risk is unpredictable, due to high variability and/or lack of power. Statistically significant increased risk of treatment failure was associated with cattle having *M. haemolytica* isolates from DNS at arrival and cattle with any positive viral PCR measured from nasal swabs, collected at first pull treatment. No other statistically significant associations with treatment outcomes were measured with any other diagnostic testing methods measured in this study, despite the power of over one thousand animals and over fourteen hundred samples analyzed.

## Discussion

Data from the current study provided evidence of poor agreement with outcome of tulathromycin treatment or metaphylaxis for BRD and bacterial culture or tulathromycin susceptibility methods used on DNS collected at arrival or at first pull from high-risk feeder heifers. Isolation of *M. haemolytica* at arrival was associated with an increased risk of BRD treatment failure; however, isolation of *M. haemolytica* at first pull treatment or isolation of *P. multocida* either at arrival or at first pull treatment was not associated with a statistically predictable risk of BRD treatment outcome. An increased RRTF associated with isolation of *M. haemolytica* on arrival does not prove causation but rather association and since viral PCR at arrival was not measured in this study, association of viral disease with isolation of *M. haemolytica* at arrival is unknown. The practicality and cost of culturing cattle at arrival becomes more economically reasonable with high prevalence of *M. haemolytica* however, the prevalence of *M. haemolytica* cannot be known without testing. Poor correlation of clinical outcomes to culture results in this study were similar in results reported by Taylor.^14^ Taylor et. al, concluded that culture results for BRD pathogens have slightly better usefulness if combined with other diagnostic methods such as clinical signs. Poor agreement of tulathromycin susceptibility testing from DNS collected at arrival and first treatment with BRD clinical outcomes, measured in this study by an experienced investigating veterinarian, provides evidence that relying on susceptibility testing to evaluate or predict tulathromycin efficacy would be unreliable.

Some investigators feel that sampling the upper respiratory tract with DNS, leads to potential sampling errors and therefore only the lower respiratory tract should be sampled.^27^ Methods to culture the lower respiratory tract in live cattle are available but not commonly used in the field due to lack of familiarization with the methods and potential risks of implementation in feed yard environments. Methods such as bronchiolar lavage, might lead to better correlation of culture results with clinical outcome in individual animals and research correlating bronchiolar lavage results to clinical outcomes is warranted. However, unless the impact of viral infections on the mucocilliary apparatus and other innate immune functions as well as presence of additional bacterial infections, e.g., *Mycoplasma bovis*, are accounted for, agreement of any diagnostic test with clinical outcome of BRD will likely remain unreliable because of the complex nature of the disease. Applying individual animal results to a population of cattle is also problematic because of the unpredictability of bacterial and/or viral pathogens involved in BRD in addition to variations in immune system function in cattle with inconsistent risk factors, i.e., risk factors and pathogens are not consistent in individual cases of BRD.^7^ **Table 7.** Bacterial culture and tulathromycin MIC results in this study involving 1026 high-risk feeder heifers and 1425 samples, indicate that the use of culture and tulathromycin susceptibility testing from DNS for making treatment decisions about the effectiveness of tulathromycin for BRD treatment or control is not trustworthy. Poor PPV and NPV, lack of statistical association of RRTF with resistant isolates and treatment outcomes far from the ≥85% parameters of susceptibility testing, measured in this study, validate this. McClary reported similar results in a retrospective study of tilmicosin susceptibility results and clinical outcome from 1297 cattle with bacterial isolates of BRD pathogens in 16 controlled clinical trials.^12^ Results of both studies, indicate extreme variability in viral and bacterial co-infections, timing and location of sample collection, and immune function of cattle with BRD, leading to poor agreement between BRD clinical outcomes and susceptibility results. Any test, specific to a single bacterial pathogen, like antimicrobial susceptibility, has a distinct disadvantage for predicting BRD clinical outcome because of the unpredictable multifactorial etiologies of BRD.

Contrary to prevailing thought, cattle in this study with *M. haemolytica* or *P. multocida* isolates susceptible to tulathromycin did ***not***have substantially greater, i.e. 85-100%, treatment success and cattle with *M. haemolytica* or *P. multocida* isolates resistant to tulathromycin did ***not***have substantially more, i.e., 85-100%, treatment failures. The lack of agreement of culture and sensitivity results with BRD clinical outcome in this study resembles results from Klement and his colleagues who reported that antimicrobial susceptibility results of common bovine mastitis pathogens poorly predicted clinical outcome.^20^

Another explanation for poor agreement of tulathromycin resistant isolates of *M. haemolytica* and *P. multocida* collected via DNS from cattle identified with clinical signs of BRD and treatment success might be the potential immunomodulatory effects of tulathromycin and other macrolide antibiotics that have been reported by several investigators.^28–31^ Poor association of tulathromycin susceptibility from isolates collected from DNS with clinical outcome could also be due to poor agreement between bacterial isolates present in the upper respiratory tract and bacterial organisms causing disease in the lower respiratory tract.^11–17^ This hypothesis needs further investigation as other researchers like Godhino,^19^ DeRosa^18^ and Capik^36^ report good correlation between isolates of the upper respiratory tract and lower respiratory tract. However, there are no CLSI standards pertaining to sample method, location, treatment versus metaphylaxis application, or timing of the disease process for BRD isolates so these parameters need to be considered when interpreting susceptibility results.

Results of this study also reinforce the idea that timing of the sample is important for interpreting the results of bacterial culture and susceptibility testing. *M. haemolytica* collected at arrival was associated with lower treatment success and isolation of *M. haemolytica* at first pull was associated with greater treatment success; however, the opposite was seen from cattle with isolates of *M. bovis* at arrival or at first pull. Other investigators have shown differences in sampling cattle at arrival and at different times of the feeding period as well as sampling cattle showing signs of BRD versus cattle not showing signs of BRD or cattle that have died due to BRD. ^14, 21, 33^ Since tulathromycin was administered at arrival, potential effect(s) of prior antimicrobial administration certainly could have impacted first pull isolates. Timing of the first pull samples in this study were at least eight days after administration of tulathromycin; however, pharmacokinetic properties of tulathromycin indicate therapeutic levels for up to fourteen days.^32^ Some of the first pull cattle may have had therapeutic levels of tulathromycin potentially confounding bacterial culture or susceptibility results, i.e., tulathromycin concentrations could have been higher than dosages calculated for CLSI breakpoints because of overlapping administration of tulathromycin. Better treatment response for first pull cattle with resistant isolates collected after arrival administration of tulathromycin may have been the result of tulathromycin therapy killing susceptible bacteria (and clearing the infection) yet leaving resistant bacteria to be cultured. Similar confounding can be expected when culturing lung tissue collected at necropsy because antimicrobial therapies administered prior to culture increase the probability of having antimicrobial levels present in tissues as well as increase the probability of culturing resistant isolates due to prior antimicrobial treatment successfully eliminating susceptible isolates.^33^ It is important to understand that although antimicrobial susceptibility testing of isolates collected from lung tissues has the advantage of sampling the site of bacterial infection, it is still inclined to confounding factors of an in vitro test for a single bacterial pathogen in a polymicrobial disease like BRD such as: sample timing relative to disease, previous antimicrobial use, additional bacterial infections, applying individual animal results to unpredictable and highly variable cattle populations and irregular influence of cattle’s immune response. Clinical outcome of treatment protocols for populations of cattle may also be confounded by prevalence of natural antimicrobial resistance in bacterial microbiomes or phenotypic expression of resistant genes following antimicrobial therapy.^33^

Another limitation of this study was the low prevalence of resistant isolates to tulathromycin in arrival cattle. The population of high-risk feeder heifers was selected to mitigate bias from variable proportions of steers and bulls and the risk of BRD associated with castration. This population was also selected because tulathromycin resistance was found in samples from previous cattle from these sources that also had poor TSR to tulathromycin, which was assumed to be from high prevalence of tulathromycin resistance. Because tulathromycin resistance at arrival was expected to be the limiting factor for sample size, the arrival and first pull susceptibility tests were combined. The authors admit that this is a limitation to the internal validity of the study because arrival and first pull cattle can be considered different populations. However, considering the low prevalence of tulathromycin resistance at arrival, in this population of high-risk feeder heifers, the authors feel, interpretation of poor agreement of tulathromycin resistance with first pull treatment failure, is still meaningful. The external validity of this study applies to the agreement of these diagnostic test methods with clinical outcome following BRD tulathromycin metaphylaxis/treatment of high-risk feeder heifers purchased in auction facilities in the southeastern U.S. or south-central Texas and transported over 8 hours to feed yard facilities in the Southern and High Plains.

In summary, potential limitations of using the results of bacterial culture and susceptibility for selecting antimicrobial agents for BRD treatment or control are: 1) bacteria isolated in the upper respiratory tract via DNS sampling may not match pathogens causing infection in the lower respiratory tract, 2) susceptibility results are specific to one bacterial pathogen and coinfections with viral or additional bacterial pathogens are common with BRD, 3) some antibiotics such as tulathromycin can have other in vivo effects besides inhibiting bacterial growth or killing bacteria such as enhancing innate immune function, and 4) the animal’s immune function may be sufficiently robust to clear infections without antibiotics or immunocompromised to the level that susceptible antimicrobials are not effective.

Multiplex PCR data collected in this study, show statistical increased risk of treatment failure with positive viral PCR swabs collected at first pull treatment. However, data from these high-risk feeder heifers, indicate negative viral PCR results collected at first pull treatment, did not reliably agree with clinical outcome. Due to the influence of disease prevalence, PCR testing at first pull is less useful with a lower prevalence of viral pathogens but more useful with a higher prevalence of viral pathogens however, this again presents the conundrum that viral pathogen prevalence is unknown without testing. A potential confounding effect of viral PCR results in this study, was vaccination with a pentavalent modified live virus vaccine at arrival and 10 to 14 days after arrival. Revaccination was not in the initial protocol but due to high morbidity, the investigating veterinarian and owner of the cattle requested an amendment to the protocol, allowing revaccination. Inability to discern vaccine virus from wild virus with a PCR test limits the diagnostic usefulness of multiplex PCR from nasal swabs collected at first pull, due to confounding from modified live virus vaccination.^33^ “Confounding does not change the fact that RRTF in this study was statistically increased with a positive viral PCR test at first pull treatment however, interpretation of those results is problematic because it is uncertain whether the viral DNA detected was from live or dead viruses or if the DNA came from vaccine or wild viruses. BRSV virus does not replicate systemically after vaccination with the SQ pentavalent MLV vaccine used and would not be expected to be recovered in the nasal passages post vaccination and therefore would not present confounding for the BRSV samples. Waltz et. al. published a trial studying the shed of vaccine virus used in this study and found no positive BRSV PCR results from DNS when sampled at 3,5,7,14,21,28,35, or 42 days post vaccination.^37^ Waltz found 20% positive at 14 days and 10% positive at 21 days for IBR and 30% positive at 7 days and 40% positive at 14 days for BVDV after vaccination with the vaccine used which would account for confounding on the BVDV and BHV-1 samples. The meaningfulness of these viral PCR results should be interpreted considering both the RRTF and prevalence, i.e., RRTF is higher for BVD and BHV-1 but prevalence was higher for BRSV and PI3V leading to similar risk of BRD treatment failure for all viral pathogens. However, further investigation is needed to more clearly define the correlation of vaccine or wild virus to better interpret results of BRD pathogen PCR testing.

Results of this study and the McClary study suggest BRD treatment failures are more likely due to additional bacterial or viral co-infections, less-than-ideal treatment timing or diagnosis, or immunocompromised animals rather than inherent bacterial resistance to the antimicrobial used for BRD treatment or control. Analysis of the data from this study suggests that relying solely on bacterial culture and tulathromycin susceptibility testing from DNS, which did not reliably agree with treatment outcome, as the primary driver in determining BRD tulathromycin metaphylaxis/treatment protocols would not be prudent. Considering the time, expertise, and expense involved in doing these diagnostic procedures and the lack of agreement of BRD clinical outcome with diagnostic test results in this study, (validated with lack of statistical risk of treatment failure, poor sensitivity, specificity, and predictive values, and treatment responses much less than 85%), the dependability of bacterial culture and tulathromycin susceptibility for evaluating the effectiveness of tulathromycin for the control or treatment of BRD appears to be unreliable.

Considering the lack of agreement with clinical outcomes and the limitations of these BRD diagnostic test methods and principles of evidence based medicine, evidence from controlled clinical trials, or meta-analysis studies would provide more reliable information for developing BRD treatment protocols and combating antimicrobial resistance compared to relying solely on diagnostic results specific to individual pathogens. Poor correlation of antimicrobial susceptibility to clinical outcome of BRD observed in this study with tulathromycin, and McClary’s study with tilmicosin, also suggests that focusing on BRD clinical efficacy data from sources with less inherent bias, to reduce the number of antimicrobial treatments, might be a more judicious way to reduce the incidence of resistant pathogens, opposed to focusing solely on pathogen MICs.

## Conclusions

While inferences to the predictive value of susceptibility testing for other antibiotics cannot be drawn, results from this study indicate that relying solely on the use of tulathromycin susceptibility testing for isolates of *M. haemolytica* and *P. multocida* derived from deep nasopharyngeal swabs collected at arrival or first pull from high-risk feeder heifers, poorly predicted clinical outcome for tulathromycin metaphylaxis/treatment.

The 95% confidence interval for RRTF measured in this study indicates that positive multiplex PCR test results from nasal swabs collected at first pull were more reliable than tulathromycin susceptibility testing for predicting clinical failures of tulathromycin for the treatment of BRD. In other words, treatment failures in these high-risk cattle, had significantly greater agreement with PCR evidence of viral involvement than resistance to tulathromycin.

Considering both this and McClary’s studies reveal poor agreement of antimicrobial susceptibility results with clinical outcome of BRD, better development of BRD treatment or control protocols might come from using information from sources such as randomized controlled clinical trials or meta-analysis studies, with better control of bias and confounding, rather than using bacterial culture and tulathromycin sensitivity results to determine treatment choices. Results of this study indicate using bacterial culture and tulathromycin susceptibility testing from DNS from high-risk feeder heifers, for predicting the efficacy of tulathromycin for metaphylaxis/treatment of BRD, is unreliable due to unpredictable or variable biases.

## Acknowledgements

Audie Waite – assistant investigator, Agri Research Center Microbial Research Incorporated, Fort Collins, CO

Texas A&M Veterinary Medical Diagnostic Laboratory, Amarillo, TX

This study was funded by Zoetis and authors except D. Bechtol (Agri Research Center) are employed by Zoetis

## Abbreviations

BA: Blood agar
BHV-1: Bovine Herpes virus type 1
BQA: Beef Quality Assurance
BRD: bovine respiratory disease
BRSV: Bovine Respiratory Syncytial virus
BVDV: Bovine Viral Diarrhea virus
CAS: clinical appearance score
CLSI: Clinical and Laboratory Standards Institute
DNS: deep nasopharyngeal swab
HFA: Hayflick’s agar
*H. somni*: *Histophilus somni*
*M. bovis*: *Mycoplasma bovis*
*M. haemolytica*: *Mannheimia haemolytica*
MHB: Mueller Hinton broth
MIC: minimum inhibitory concentration
Multiplex PCR: BVDV, BRSV, BHV-1, PI-3V
NPV: Negative Predictive Value
PCR: polymerase chain reaction
PI-3V: Bovine Parainfluenza virus type 3
PMI: Post Metaphylaxis Interval
*P. multocida*: *Pasteurella multocida*
PPV: Positive Predictive Value
PTI: Post Treatment Interval
RRTF: relative risk for treatment failure
TFR: treatment failure rate
TSR: treatment success rate

Research locations: Animal research - Agri Research Center, Canyon, Texas

Laboratory research - Microbial Research Incorporated, Fort Collins, Colorado

Texas A&M Veterinary Medical Diagnostic Laboratory, Amarillo, Texas Supported by a grant from Zoetis

Tulathromycin susceptibility results presented at:

Academy of Veterinary Consultants, Kansas City, KS, December 2015 Summary presentation at:

World Buiatrics Congress, Sapporo, Japan, August 2018 Poster presentation at:

BRD Symposium, Denver, CO, December 2019

Zoetis is the manufacturer of tulathromycin, Draxxin^®^, and all authors except Dr. Bechtol are employed by Zoetis

No extra-label antimicrobial use

*N/A = not analyzed due to one category with zero, i.e., treatment failure/success

